# Human decision-makers terminate evidence accumulation using flexible decision rules

**DOI:** 10.64898/2026.03.18.712662

**Authors:** I Kalburge, A Dallstream, K Josić, ZP Kilpatrick, L Ding, JI Gold

## Abstract

Decisions based on evidence accumulated over time require rules governing when to end the accumulation process and commit to a choice. These rules control inherent trade-offs between decision speed and accuracy, which require careful balance to maximize quantities that depend on both like reward rate. We previously showed that, to maximize reward rate, normative decision rules adapt to changing task conditions (Barendregt et al., 2022). Here we used a novel task to examine whether and how people use adaptive rules for individual decisions under a variety of conditions, including changes in decision outcomes across trials and changes in evidence quality both across and within trials. We found that the participants tended to use rules that adjusted, at least partially, to predictable changes in task conditions to improve reward rate, consistent with a rationally bounded implementation of normative principles. These findings help inform our understanding of the extent and limits of flexible decision formation in the brain.

## Introduction

Many “sequential-sampling” models of decision-making, like the drift-diffusion model (DDM), assume that the decision process terminates when accumulated evidence reaches a fixed, predefined threshold, or decision bound (Laming, 1968; Link, 1992; Gold and Shadlen, 2007; Ratcliff et al., 2016). The height of the bound balances an inherent trade-off between decision speed and accuracy, with lower bounds emphasizing speed and higher bounds emphasizing accuracy (Gold and Shadlen, 2002; Simen et al., 2009; Heitz, 2014). During the accumulation process, this bound is typically assumed to be either: 1) “fixed”, which under particular, static task conditions can optimize quantities like average accuracy (or reward) per unit time (Barnard, 1946; Wald, 1947; Bogacz et al., 2006) and has been used extensively to model human and animal behavior (Zhang, 2012; Heitz, 2014); or 2) “collapsing” over time, which can also be part of an optimal decision strategy under more complex and/or dynamic conditions (Malhotra et al., 2018) and has been used to model multi-choice decisions (Churchland et al., 2008), mixed task demands (Palestro et al., 2018), decisions with strong sequential effects (Urai et al., 2017; Nguyen et al., 2019), urgency (Cisek et al., 2009; Thura et al., 2012), evidence calibration (Hanks et al., 2011), and accumulation costs (Drugowitsch et al., 2012).

However, our understanding of the decision rules people employ remains incomplete, reflecting two key limitations of existing approaches to study them. First, because models like the DDM require many trials to estimate parameters from behavioral data, model variants with fixed or collapsing bounds primarily characterize trial-averaged decision rules. These approaches thus offer limited insights into the rules governing individual decisions. Second, existing models typically assume that decision-relevant quantities do not change during the deliberative process within a trial. Under such static conditions, decision rules with a fixed temporal structure (e.g., constant or governed by a predefined collapsing trajectory) can be effective. In contrast, when the decision-relevant quantities change during deliberation, flexible adjustments in evidence accumulation and/or decision rules can improve performance (Drugowitsch et al., 2012; Thura et al., 2012; Ossmy et al., 2013; Cheadle et al., 2014; Glaze et al., 2015; Hawkins et al., 2015; Tajima et al., 2016; Veliz-Cuba et al., 2016; Piet et al., 2018; Li et al., 2019; Jang et al., 2021; Barendregt et al., 2022). However, the exact nature of these adjustments, and how they operate under a broad range of conditions involving changes in task demands across and within individual decisions, are not well understood.

To address these issues, we developed and used a task that allowed us to measure the rules people use to terminate evidence accumulation in a simple two-alternative setting (the “pigeon task”; Fig. 1). The task has two important features. First, the decision variable, which is typically computed internally by the decision-maker and thus hidden from the experimenter, is presented explicitly as a dynamic visual stimulus (a pigeon following a “random walk” towards one of two piles of seeds). This design minimizes the uncertainty that both the decision-maker and the experimenter may have about the exact values and trajectory of the subjective decision variable on each trial, allowing us to focus on understanding the decision rule used to commit to a decision. Second, we measure the decision rule for each decision directly, in contrast to the conventional practice of inferring the decision rule by fitting data from many trials. We used several variants of this task to characterize how people implement these rules to make individual decisions under different conditions. The results provide new, empirical perspectives on the extent and limits of the flexibility with which we balance the speed and accuracy of decisions in changing environments.

**Figure 1:**
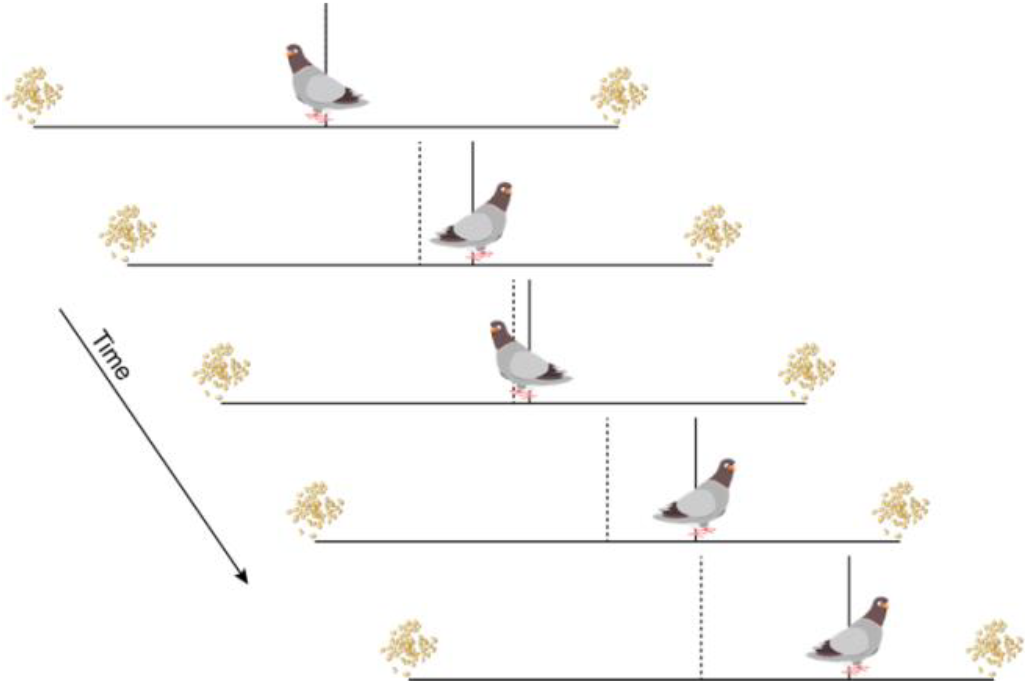
Pigeon task. On each trial, a pigeon takes a random walk towards one of two seed piles. The participant presses a key at any time to terminate the walk and indicate which pile they think the pigeon will eventually reach. Deciding quickly saves steps (which are limited to 600 per block) at the expense of accuracy. Waiting improves accuracy (which governs coins earned and lost) at the expense of more steps. Continuous feedback is given about total coins earned and steps taken. Vertical bars indicate the midpoint of the path (dotted) and the midpoint of the pigeon (solid), to allow the participant to identify their locations precisely.

## Results

We used the on-line testing platform Prolific (prolific.com) to collect behavioral data on the pigeon task. We used three versions of the task, which differed in terms of manipulations of decision outcomes (rewards and costs), the signal-to-noise ratio (SNR) of the evidence across decisions, and the SNR of the evidence within decisions, that were each performed by a separate cohort of 60 participants (see Table 1). Human protocols were approved and determined to be Exempt by the University of Pennsylvania Internal Review Board (IRB protocol 844474). Participants provided consent on-line before they began the task.

**Table 1:**
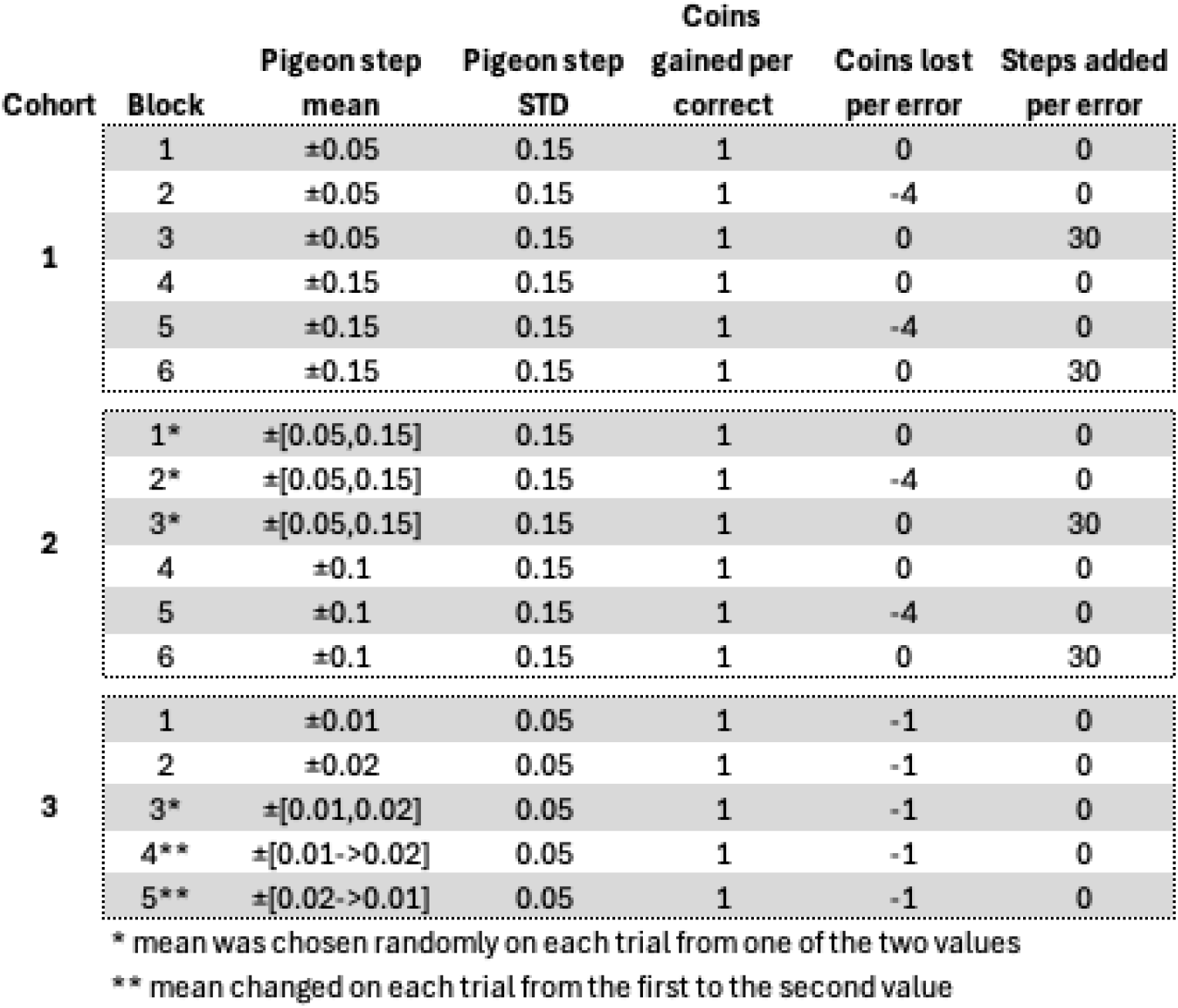
Taskparameters.

### People use decision rules representing a bound on accumulated evidence, as in many decision models

The pigeon task allowed us to observe directly, at the time of decision commitment (a keypress indicating a left or right choice), the value of the decision variable (pigeon position) that was presented explicitly on each trial. Figure 2A shows an example of pigeon trajectories and choices from individual trials for a single participant. The participants tended to choose the same side as the final pigeon position (“congruent” choices, corresponding to black symbols with positive ordinate values, gray symbols with negative ordinate values in Fig. 2A) on most, but not all, trials. We assumed that they intended to make congruent choices but occasionally made incongruent choices that resulted, in part, from random pigeon movements during an inevitable and potentially variable motor delay between mentally committing to a decision and initiating the keypress. We estimated this delay (non-decision time, or NDT) for each participant and task block by finding the delay (between 0 and 4 pigeon steps presented at 5 steps/sec) that maximized congruence (Fig. 2B), using a single value per participant for simplicity. Subtracting this participant- and block-specific non-decision time from the response time (RT) yielded the decision time (DT) on each trial. Performance was variable across participants but, as expected for a DDM-like decision process, exhibited higher accuracy with longer average DT (Spearman’s correlation between accuracy and median DT across participants, *ρ*= 0.77, *H*_*0*_: *ρ*=0, *p*<0.001 for the data shown in Fig. 2C).

**Figure 2:**
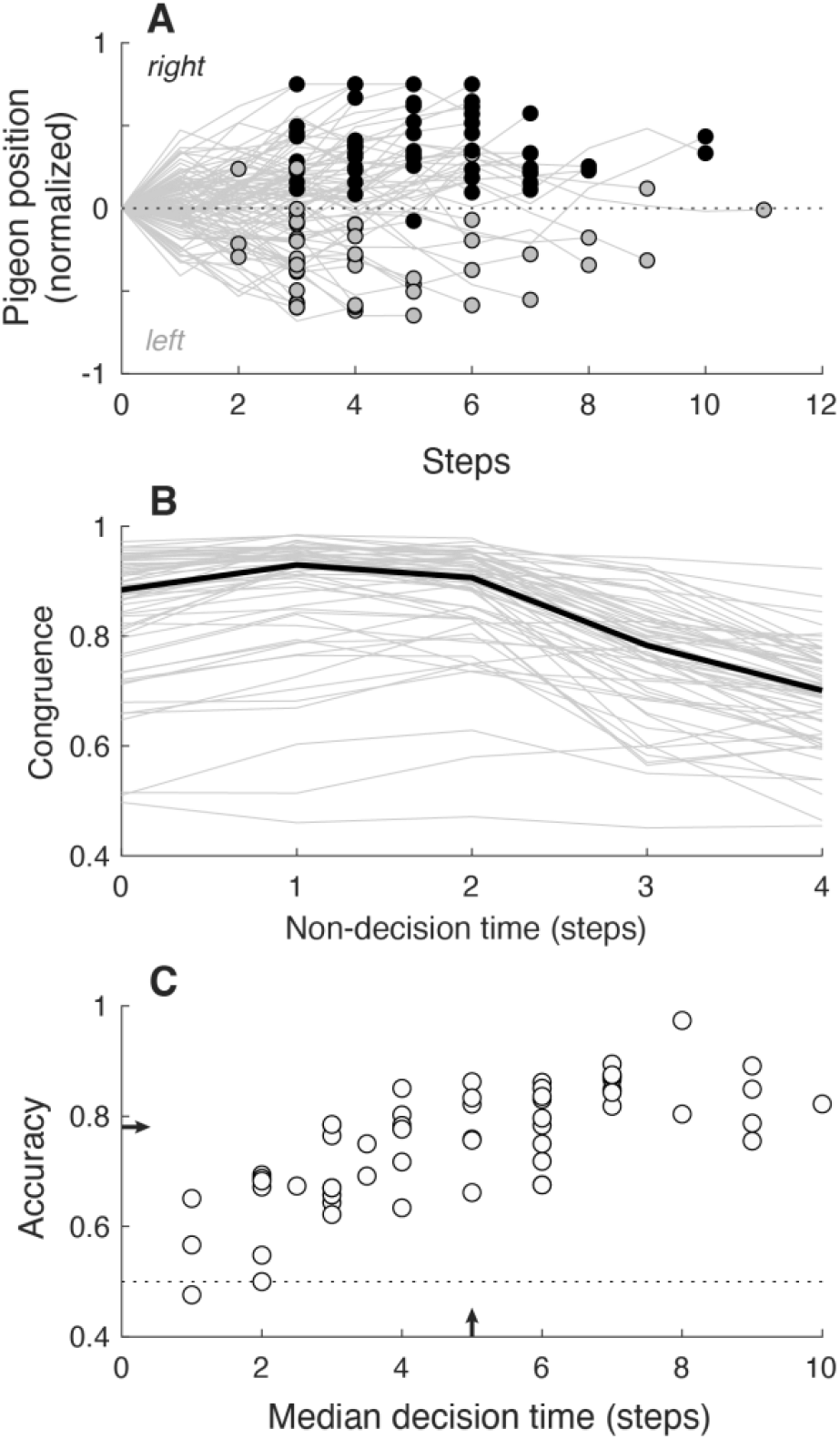
Performance summary. A. Pigeon positions (lines) and the participant’s choices (circles at the final positions; gray for left, black for right) from individual trials from an example session. B. Estimates of non-decision times (NDTs). We assumed that, except for occasional lapses, the participants tried to make choices that were congruent with the position of the pigeon at the time of the choice; i.e., they chose right/left when the pigeon was to the right/left side of center. We therefore estimated the NDT for each participant (gray lines) for a given block as the delay corresponding to the maximum value of this congruence, which tended to peak at 1–2 steps (the median congruences across participants is shown as the thick black line). DT was computed on each trial by subtracting this estimate of NDT from the measured response time (RT, in steps of the decision variable). C. Overall percent correct as a function of the median DT for each participant (individual symbols). Arrows indicate median values. Data in all panels are from cohort 1, block 2 (see Table 1).

For each trial, we defined the bound as the pigeon position at the DT. To illustrate basic properties of the bounds used by the participants, we focus first on data from cohort 1, block 2 (fixed low SNR throughout the block and a relatively large coin penalty for wrong choices; see Table 1 in Methods). We consider bound magnitude and thus, unless otherwise noted, use the absolute distance from the initial center point, normalized by the distance to the edge of the screen. The measurements did not differ between choices (Wilcoxon signed-rank test for *H*_*0*_: equal median bounds for left versus right choices, *p*<0.01 for just one of the 60 sessions, none after correcting for multiple comparisons) and were thus combined for both choices in the remaining analyses.

Under these conditions, the participants used bounds that exhibited several trends (data from an example participant are shown in Fig. 3A; data from the full cohort of 60 participants are summarized in Fig. 3B). First, the bounds appeared to be relatively low for decisions that took only ∼1–2 steps, reflecting fast guesses (instead of bounded evidence accumulation) made by many of the participants on a subset of trials. Second, at longer DTs, the bounds tended to remain relatively constant, albeit with a slight downward trend (“collapse”), on average, as a function of DT (median [IQR] slope of a linear regression of absolute bound versus DT for DT>2 = -0.01 [-0.01 0.00] z-score/step, *H*_*0*_: median=0, *p*=0.01; Fig. 3C). Third, the bounds tended to vary noisily around a relatively constant mean value across trials within a block, partly reflecting uncertainty in our ability to measure the bound because of the discrete pigeon steps (see Methods). A signature of such noisy variation is that the absolute bound change from one trial to the next should depend linearly on the bound value from the first of those two trials with a slope of one (i.e., a regression to a constant mean); consistent with this idea, the participants’ data had a median [IQR] slope of 0.92 [0.75 1.02] (Fig. 3D). Moreover, the distributions of their trial-by-trial bound choices were comparable to those obtained from simulated behavior that used the same generative statistics for the pigeon trajectories as in the experiments and assumed that the decision-maker used the same, fixed bound on each trial (Wilcoxon signed-rank test for *H*_*0*_: equal median standard deviations of bound estimates from empirical versus simulated data, *p*=0.34). These results suggest that the participants used relatively constant mean bound values, and the variability reflected primarily estimation noise.

**Figure 3:**
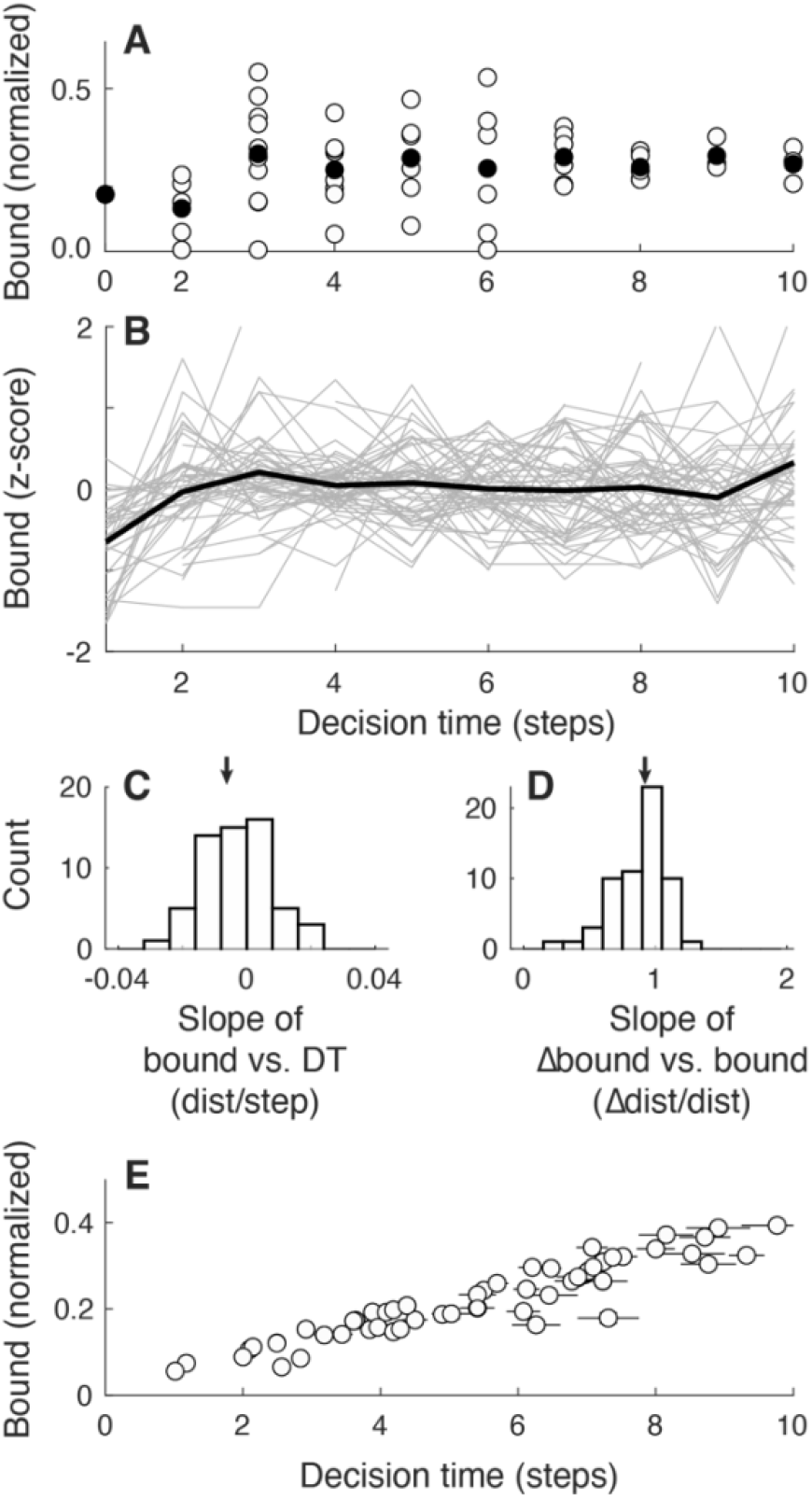
Bound summary. A. Data from one example session showing the decision bound (absolute distance of the pigeon from the central starting point assessed at the DT, normalized by the distance from the starting point to the edge of the screen) as a function of DT. Open symbols are data from individual trials. Filled symbols are medians per DT. B. Summary of data from 60 participants. Gray lines are median bounds (z-scored across all trials for each participant, to facilitate comparing time courses across participants) per DT as a function of DT for each participant. Black line is the median of the gray lines, computed per DT. Note that bound estimates are the least reliable at the shortest DTs (see Methods). C. Histogram of slopes of linear regressions of absolute bound per DT, computed separately for each participant (excluding DT<=2 to avoid unreliable bound estimates at the shortest DTs). Values ∼0 imply no systematic (linear) dependence on DT. D. Histogram of slopes of linear regressions of the change in bound magnitude versus the previous bound, computed separately for each participant. Values ∼1 imply that trial-to-trial bound adjustments were consistent with a regression to the mean (i.e., noise). Arrows in C and D indicate median values. E. Mean±sem bound height (symbols and vertical error bars) as a function of mean±sem DT (circles and horizontal error bars) computed per participant. Data in all panels are from cohort 1, block 2 (see Table 1).

In addition to these within-participant trends, across participants there was a strong, positive relationship between average DT and average bound per participant (Spearman’s *ρ*=0.95, *H*_*0*_: *ρ*=0, *p*<0.001; Fig. 3E), consistent with the positive relationship between DT and accuracy (Fig. 2C). Together, these results imply that, under these fixed conditions, people tended to use decision rules that were consistent with predefined, relatively fixed bounds like in standard accumulate-to-bound models.

### People use decision rules that adjust to changes in decision outcomes that affect reward rate

The standard accumulate-to-bound decision process involves inherent trade-offs between speed and accuracy that, in principle, can be optimized with respect to quantities that depend on both like reward rate (Gold and Shadlen, 2002). Previous work has shown that certain well-performing participants use bounds that conform to optimal predictions (Simen et al., 2009). Here we examined in more detail how the decision bounds used by individuals relate to reward-rate functions across a variety of manipulations of the rewards and costs associated with decision outcomes (cohort 1, blocks 1–3; see Table 1). Specifically, each of three blocks consisted of a fixed number (600) of total pigeon steps that were decremented on each trial by one initial step to start the trial plus the number of steps taken until the keypress (i.e., the RT). Correct responses were associated with a reward (coins gained), and error responses were associated with a cost, either in coins or steps lost. Because we used a fixed number of steps, reward rate was equivalent to the total number of coins earned within a block. After completing a session, each participant was given a bonus payment proportional to the total number of coins they earned across all blocks.

Under these conditions, a fixed-bound DDM makes the following predictions about the dependence of expected reward rate on bound values (Fig. 4A–C, red lines). For block 1 (+1 coin gained for a correct response, 0 coins or steps lost for an error), relatively low bounds yield the highest reward rates because errors are not punished and thus fast guessing is an effective strategy. For block 2 (+1 coin gain for a correct response, -4 coins lost for an error), low bounds yield the lowest reward rates, because errors are punished with a coin cost and thus gathering more evidence to improve the accuracy of each choice is an effective strategy. Reward rates increase rapidly as bounds increase and then plateau and reach a peak for bound values ∼0.4. For block 3 (+1 coin gain for a correct response, -30 steps lost for an error), reward rate is affected minimally by bound height, because this cost in steps discouraged guessing but did not provide a substantial advantage for any particular level of accuracy. These relationships predict that participants sensitive to reward rate should increase their bounds from block 1 to block 2 but make minimal adjustments from block 2 to block 3.

**Figure 4:**
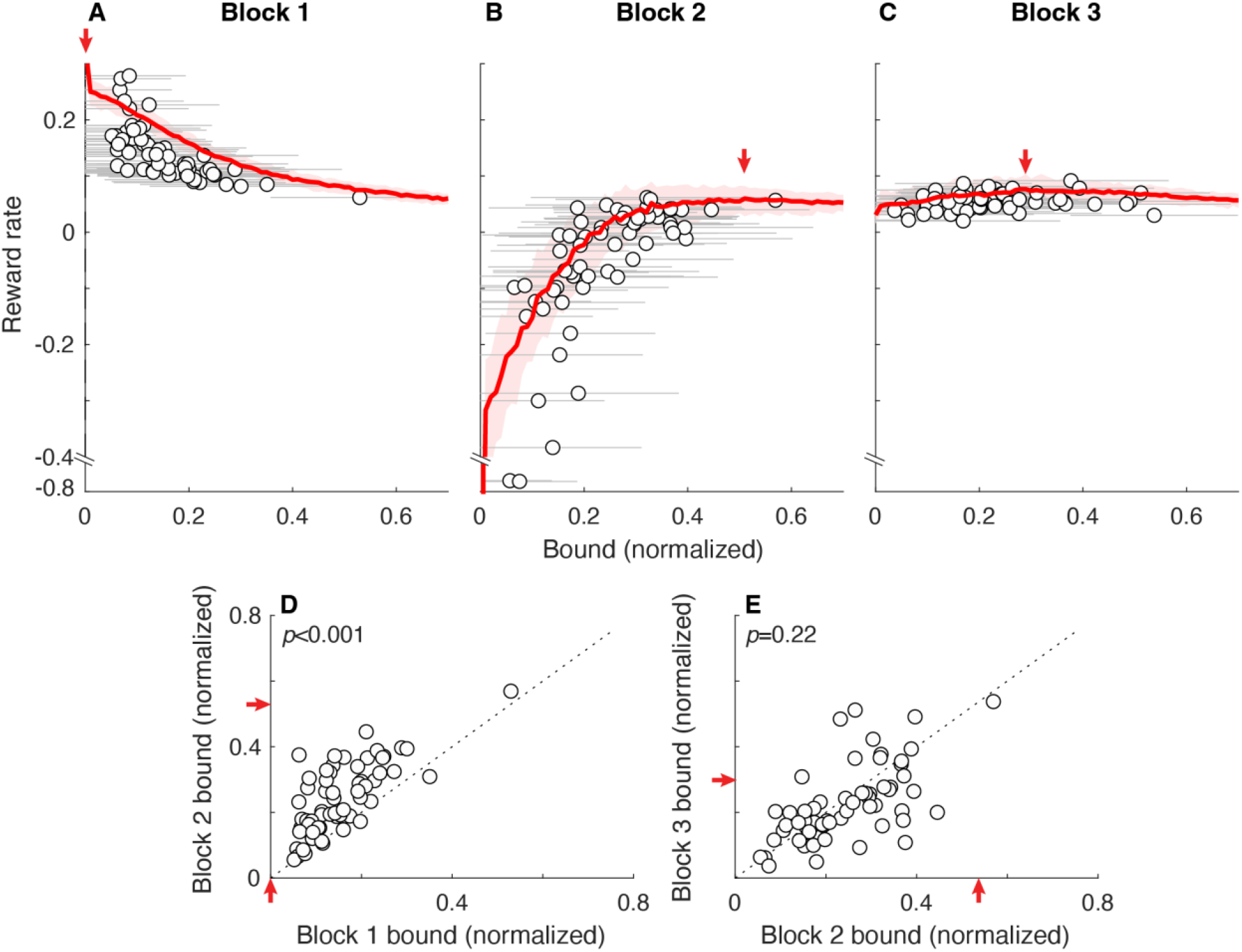
Block-wise bound adjustments to changes in rewards and costs. 60 participants each performed in sequence three blocks of trials (each block ended after 600 total pigeon steps) that differed in terms of the reward and cost structure: A) block 1, +1/0 coins for correct/error responses, no step cost; B) block 2, +1/-4 coins, no step cost; and C) block 3, +1/0 coins, –30 steps for an error. Each panel shows reward rate (coins/step; note the broken axis to include the two outlier participants in block 2) as a function of absolute bound. The red lines and ribbons are the medians and 95% CIs from simulations using the given fixed bound. Points and error bars are mean±SEM bounds and experienced reward rate from individual participants. The observed spread in reward rates around the model prediction is largely attributable to trial-by-trial variability in participant bound usage. D, E. Comparison of bounds between block 1 and 2 (D) and between block 2 and 3 (E) for individual participants (points). Arrows in all panels point to reward-rate maxima. *P*-values are for Wilcoxon signed-rank test for *H*_*0*_: median difference between *x* and *y* values=0.

Consistent with these predictions, the participants tended to make systematic, upward adjustments of their decision bounds from block 1 to block 2, corresponding to an increase in the cost of making errors (Fig. 4D; median [IQR] difference=0.08 [0.04 0.13], Wilcoxon signed-rank test for *H*_*0*_: median difference=0, *p*<0.001). These adjustments were consistent with a sensitivity to reward-rate functions that peaked at lower bounds (emphasizing speed) for block 1 and higher bounds (emphasizing accuracy, plateauing across a fairly large range of values after a steep increase at lower bound values) for block 2. In contrast, the participants did not make systematic adjustments to their decision bounds from block 2 to block 3 (median [IQR] difference=-0.03 [-0.06 0.03], *p*=0.22; Fig. 4E). This lack of adjustments was consistent with a relatively flat reward-rate function in block 3, which implied a minimal marginal benefit for any change in decision bound.

The participants also appeared to maintain individualized tendencies in their bound choices across blocks. Specifically, their bounds varied considerably across individuals but were relatively stable across blocks within individuals (Spearman’s ρ comparing block 1 to block 2=0.68, *H*_*0*_: *ρ*=0, *p*<0.001; block 2 to block 3=0.63, *p*<0.001). That is, participants who adopted relatively low bounds in one block tended to remain low relative to others across blocks, and likewise for high-bound participants. Together these results indicate that the decision bounds used by the participants were not chosen at random but instead reflected a combination of internal and external factors that included a sensitivity to rationally bounded (good-enough) reward rates.

### Decision rules adjust to expected, but not unexpected, across-decision differences in evidence quality

Reward rates are sensitive to not just outcome-related costs and benefits but also the strength of the evidence used to form the decision. Understanding how people account for this sensitivity to evidence strength is highly relevant to many studies of decision-making that involve experimentally controlled changes in evidence strength. When these changes occur predictably across blocks, it is typically assumed that decision-makers adjust their decision processes (including scaling of the evidence and/or bounds) by evidence strength to optimize performance. In contrast, when these changes in evidence strength occur unpredictably from trial to trial, it is typically assumed that decision-makers do not first infer evidence strength and then use that inference to adjust decision parameters within each trial. These assumptions tend to be used in decision models that capture trial-averaged human decision-making behavior (Bogacz et al., 2006; Shadlen et al., 2006). Here we examined in more detail if and how individual decision-makers adjust their decision bounds in response to predictable (block-wise) and unpredictable (trial-wise) changes in evidence strength (here measured as the SNR of the decision variable).

When the SNR of the decision variable was varied across blocks, the participants tended to make adaptive bound adjustments in the direction of maximizing reward rate (Fig. 5A,B). Under these conditions, a fixed-bound DDM can increase the overall reward rate by using a higher bound for the low-versus high-SNR block (heatmap in panel A). This difference occurs because a higher bound is needed to provide the same weight of evidence (i.e., a quantity that scales with the logarithm of the likelihood ratio to support optimal decision-making) for the low-versus high-SNR condition (Gold and Shadlen, 2001). The participants’ behavior was, on average, consistent with this optimal strategy: average bounds tended to be higher in the low-SNR block than in the high-SNR block (cohort 1, blocks 2 and 5; Wilcoxon signed-rank test for *H*_*0*_: equal medians, *p*<0.001; Fig. 5A). Crucially, this difference in bounds was evident across a range of decision times (Fig. 5B). These results are not consistent with SNR-independent collapsing bounds alone, which predict lower bounds for low-SNR blocks (because low SNR corresponds to longer decision times and thus a higher probability of reaching a more collapsed bound) and no difference in bounds on high-versus low-SNR trials measured for the same decision time.

**Figure 5:**
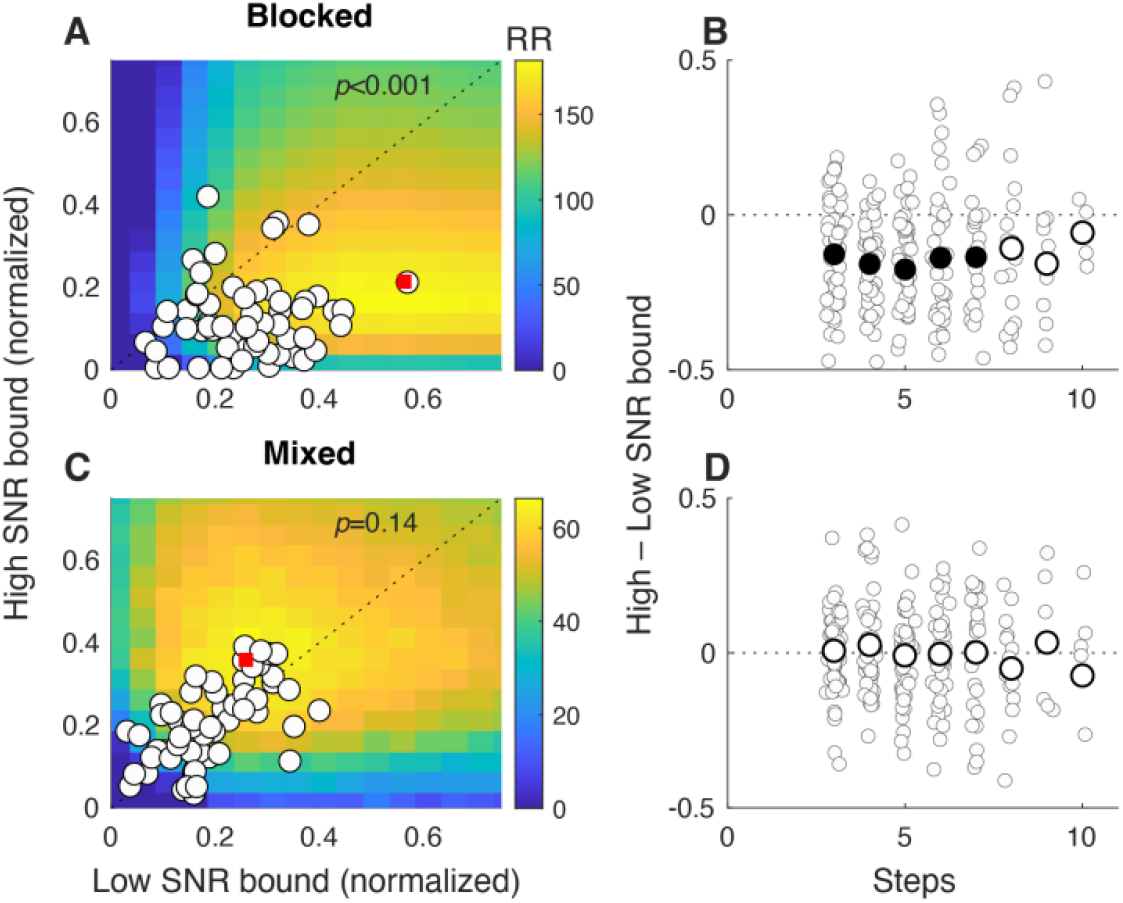
SNR-dependent bounds. A,C. Bounds from high-SNR (ordinate) versus low-SNR (abscissa) trials, when SNR changed by block (A) or trial (C). Heatmaps show the predicted reward rate (RR, colorbars to the right; maxima indicated by red squares) for different combinations of bound values for high and low SNR blocks (A) and trials (C). Circles denote median SNR-specific bounds from individual participants, which were systematically higher for low-versus high-SNR block-wise changes (A) but not reliably separated for trial-wise changes (C). *P*-values are for Wilcoxon signed-rank test for *H*_*0*_: median difference between *x* and *y* values=0. B, D. Difference in median bound between high-SNR and low-SNR trials as a function of decision time (DT). Small circles indicate data from individual participants, large circles are medians across participants, per DT. Filled large circles indicate *p*<0.05 for *H*_*0*_: median=0.

In contrast, when the SNR was chosen randomly (high or low) on each trial in a block, participants tended to use the same bound for each SNR condition (cohort 2, block 2; Fig. 5C,D). Under these conditions, a decision-maker using different, fixed bounds for the two SNRs (e.g., by quickly perceiving the SNR difference and setting the bound accordingly) would maximize overall reward rate by using a higher bound for the high-versus low-SNR trials (brighter yellow colors in panel C). This difference occurs because with a fixed number of steps per block, it is better to use those steps to emphasize accuracy on high-SNR trials and thus maximize overall reward rate. However, the participants did not tend to follow this prescription and instead used similar, average bounds on both high- and low-SNR trials and across a broad range of decision times (Wilcoxon signed-rank test for *H*_*0*_: equal medians, *p*=0.14; Fig. 5C,D). These participants also tended to use similar, minimally collapsing bounds when SNR was mixed (block 2) or fixed (block 5) across trials (median slopes = -0.002 and -0.012 bound change/step for mixed and fixed, respectively; Wilcoxon signed-rank test for *H*_*0*_: median difference of slopes of linear regression of bound versus decision time for mixed versus fixed SNR = 0, *p*=0.10). Thus, the participants were generally able to adapt their bounds to predictable blockwise changes, but not unpredictable trialwise changes, in evidence quality.

### Decision rules adjust to expected, within-decision changes in evidence quality

A novel prediction of our previous work is that in certain dynamic environments, normative decision rules that are used to maximize reward rate can include changes in bounds that occur not just across, but also within, individual decisions (Barendregt et al., 2022). Among the simplest conditions in which these adjustments are expected to occur is when there is a single, predictable changepoint in evidence quality (SNR) during decision formation. We tested this idea on a third set of participants (cohort 3). Each participant first performed versions of the task with no changepoints and either a (relatively) high or low SNR (we chose SNR values with substantial effects on expected reward-rate functions when SNR changed within a trial, but were similar enough to have minimal effects on reward-rate functions when SNR was held steady within and across trials but varied across blocks; see Fig. 6F heatmap). They then performed versions with changepoints, both low→high SNR (in which the timing of the changepoint was set to the median RT from the low-SNR, no-changepoint condition; median [IQR] changepoint times = 9.5 [8.0 15.0] steps) and high→low SNR (in which the timing of the changepoint was set to the median RT from the high-SNR, no-changepoint condition; median [IQR] changepoint times = 8.0 [6.5 11.0] steps).

**Figure 6:**
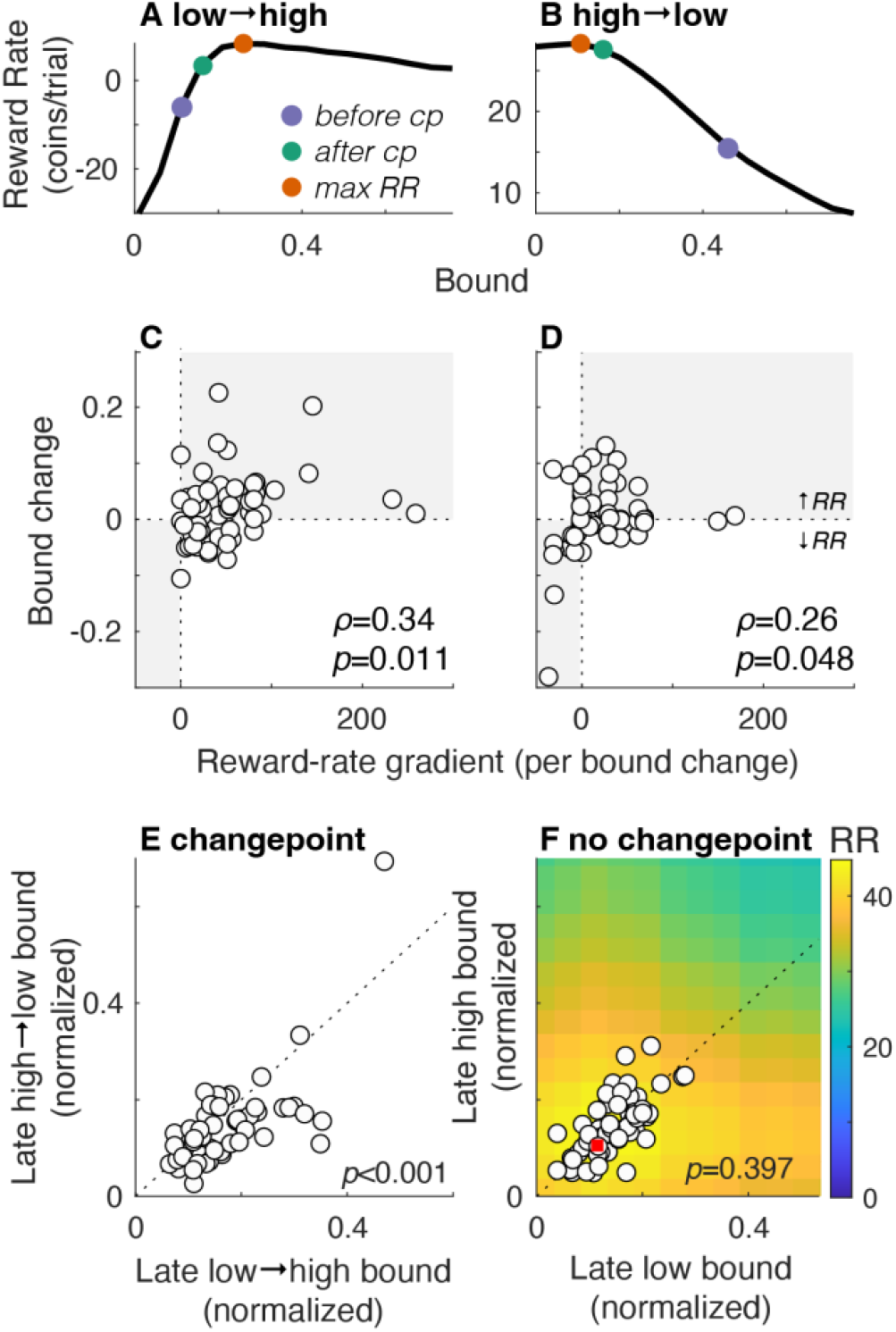
Changepoint-dependent bounds. A,B. Reward rate functions of post-changepoint bound height depend on the time of the changepoint and the bound used before the changepoint. Example functions from one participant performing the low→high task (A) and one performing the high→low task (B). Red points indicate the peaks of each function. The other points indicate the average bound used by each participant before (blue) and after (green) the changepoint. To increase reward rate, the participant in A increased their bound after the changepoint, whereas the participant in B decreased their bound after the changepoint. C,D. Change in bound from pre-to post-changepoint decisions (ordinate) versus reward-rate gradient (change in reward per change in bound to achieve the maximum reward rate; positive/negative values indicate that reward rate increases by increasing/decreasing the bound after the changepoint) computed for each participant (circles) for low→high (C) and high→low (D) SNR conditions. Changes that increase reward rate are in the gray quadrants. Spearman’s *ρ* and associated *p*-values are shown in each panel. E. Median bound for post-changepoint decisions for high→low (ordinate) versus low→high (abscissa) SNR conditions for individual participants (circles). F. Median bound for late decisions (i.e., after the median RT) for high (ordinate) versus low (abscissa) SNR conditions for individual participants (circles). Heatmap indicates expected reward rate (RR, colorbars to the right; maximum is indicated with a red square). *P*-values in E and F are for Wilcoxon signed-rank test for *H*_*0*_: median difference between *x* and *y* values=0.

Under these conditions, maximizing reward rate requires using a decision bound that changes within a trial (Barendregt et al., 2022). The details of the dynamics depend on, and are generally congruent with, the direction of the change in SNR. For low→high SNR changes, the normative bound should increase, on average, at the changepoint, because after waiting past the changepoint is worthwhile to accumulate more of the stronger evidence to improve accuracy. In contrast, for high→low SNR changes, the normative predictions for bound changes at the changepoint are more complicated because they depend strongly on the bound used before the changepoint, which varied considerably across participants. When the pre-changepoint bound is low, DTs tend to be very short, and thus raising the bound at the changepoint tends to improve reward rate by observing a few extra steps to increase accuracy. In contrast, when the pre-changepoint bound is high, the bound should be lowered at the changepoint to improve reward rate by not wasting time with extra, low-SNR evidence.

To account for these variable dynamics, we computed the dependence of expected reward rate on post-changepoint bound for each participant. We considered both the time of the changepoint (determined for each participant from their RTs on a preceding non-changepoint block) and the average bound used on trials that ended before the changepoint (i.e., their “initial bound”), computed separately for each participant. Two example reward-rate functions are shown in Fig. 6A,B: for the example in panel A, the participant increased their bound to increase reward rate, whereas for the example in panel B, the participant decreased their bound to increase reward rate.

Overall, the participants’ behavior was roughly consistent with these normative changes. For each participant, we compared their actual average bound change at the SNR changepoint (the difference in the mean bound for trials that ended before versus after the changepoint; positive/negative values imply bound increases/decreases) to the reward-rate gradient from their pre-changepoint bound to the bound associated with the maximum reward rate (i.e., how much reward rate is expected to change per unit bound change in the direction of maximum reward rate; positive/negative values imply bounds should increase/decrease to maximize reward rate). These values were positively correlated for both the low→high (Spearman’s ρ=0.34, *H*_*0*_: ρ=0, *p*=0.011; Fig. 6C) and the high→low (ρ=0.26, *p*=0.048; Fig. 6D) SNR condition. These bound changes corresponded to consistently larger post-changepoint values of bound height in the low→high than the high→low condition (Wilcoxon paired signed-rank test for *H*_*0*_: equal median change, *p*=0.006; Fig. 6E). These bound differences were not simply a result of SNR-dependent differences in bounds occurring late in a trial, independent of the changepoint: the same participants, performing blocks of non-changepoint trials, used bounds on trials in which the choice occurred after their median RTs that were not statistically distinguishable on low-versus high-SNR trials, as expected from the reward-rate function (*p*=0.397; Fig. 6F; note that this comparison differed from Fig. 5A in that only long-DT trials were included here).

We also verified that the initial bounds used by participants were similar for the same initial SNRs, regardless of the presence or absence of a changepoint; i.e., between low-SNR fixed blocks and low→high changepoint blocks (Wilcoxon paired signed-rank test for *H*_*0*_: equal medians, *p*=0.723) and between high-SNR and high→low changepoint blocks (*p*=0.098). Thus, the bound adjustments on changepoint trials did not include changes in initial bound values in anticipation of a changepoint but rather occurred within each trial around the (predictable) time of the changepoint.

## Discussion

Deliberative decisions free the brain from the immediacy of reflexive processing but pose a critical challenge: how does the brain decide when to stop deliberating and commit to a proposition or course of action? Our understanding of this commitment process has been dominated by a computational framework that assumes that decisions are terminated once the accumulated evidence reaches a particular level, or bound, that was fixed at that level before any evidence was obtained. This kind of rule has close ties to normative theory and can explain a range of behavioral and neural findings but is optimal only under the highly restrictive conditions used in many decision studies: when the informativeness, rate of acquisition, and other features of the evidence are stable and known in advance (Laming, 1968; Link, 1992; Gold and Shadlen, 2007; Heitz, 2014; Ratcliff et al., 2016). Guided by previous findings identifying exceptions to this kind of fixed bound, including bounds that “collapse” over time (Hawkins et al., 2015; Voskuilen et al., 2016; Palestro et al., 2018) and our recent theoretical work identifying environments in which adaptive, time-dependent bounds are optimal (Barendregt et al., 2022), we sought a more detailed understanding of the kinds of decision rules people use under varied and dynamic conditions.

Our novel task design enabled direct measurement of the decision bound at the time of commitment on each trial. Consistent with many “accumulate-to-bound” models, we found that participants tended to commit once the decision variable reached a characteristic level, with modest collapse over time and trial-to-trial variability. Subjective bound heights, which mediate the trade-off between speed (lower bounds) and accuracy (higher bounds), varied considerably across individuals but for each individual were relatively stable across task conditions, representing different speed-accuracy preferences that can arise from a variety of factors (Ho et al., 2012; Perri et al., 2014; Fiedler et al., 2021). In addition, the participants tended to adjust their bounds in directions that improved, but did not necessarily optimize, reward rate. That is, bound heights clustered around “good enough” regions of the reward rate function, suggesting a bias towards “satisficing”, rather than strictly reward maximizing, strategies, especially when the reward-rate function was shallow near its maximum.

We also present three findings that relate decision bounds to the signal-to-noise ratio (SNR) of the decision variable. First, changing SNR across blocks, but keeping it fixed within blocks and trials, led to systematic changes in decision bounds, with generally higher bounds for low-versus high-SNR blocks. These adaptations were consistent with normative theory, which assigns a stronger weight-of-evidence to higher-SNR signals (Bogacz et al., 2006; Tardiff et al., 2025) that thus require lower bounds. These results also demonstrated that participants tended to choose decision bounds in satisficing reward-rate regimes.

Second, changing SNR across, but not within, trials, did not have systematic effects on decision bounds. Under these conditions, which are typical for many perceptual tasks, it is often assumed that the decision-maker sets the bound on that trial before assessing the SNR online. Accordingly, behavioral data are often well described by models that assume the same bound is used across trials. In some cases, these models include a collapse over time that has been proposed to be a surrogate for SNR-dependent adjustments, because the longer a trial has gone on without the bound being crossed, the lower the SNR is likely to be and thus the lower the normative bound should be (Hanks et al., 2011). Our participants did not tend to use collapsing bounds under these conditions but instead adopted a single bound closer to the high-SNR regime, consistent with preserving speed on easy trials rather than protecting accuracy on difficult trials. These results are consistent with previous findings that the specific form of bound under these kinds of conditions can depend strongly on the exact task conditions (Hawkins et al., 2015; Palestro et al., 2018).

Third, changing SNR at a predictable time and in a predictable direction within a trial caused systematic adjustments in decision bounds after the SNR changepoint. Our task design makes these effects directly observable on a trial-by-trial basis, allowing decision bounds to be measured with minimal reliance on trial averaging or model-based inference. This general result was predicted by our recent theoretical work (Barendregt et al., 2022), albeit with some key caveats. Like the theory, our results showed that bounds tended to increase or decrease with within-trial increases or decreases, respectively, in SNR. However, our results also included across-participant differences in the timing of the changepoint (to ensure that approximately half of the trials ended before and half ended after the changepoint) and the bound height used before the changepoint (which varied considerably across, but not within, participants). Both factors influence the relationship between post-changepoint bound height and expected reward rate, including more complicated dynamics when SNR changes from high to low.

Together these results imply decision rules are governed by both (near-) optimality and predictability. As we noted, the participants tended to use bounds that achieved satisficing, but not truly optimal, reward rates. Reward rate is a convenient metric to use in our analyses, but it seems likely that people operate under additional constraints (e.g., in computational and other resources; Simon, 1972). The local steepness of the reward-rate function near its maximum likely determines how strongly behavior is driven to fine-tune decision bounds: when the peak is shallow, a wide range of bounds yields nearly equivalent reward rates, reducing the incentive or pressure to adapt precisely (e.g., compare Figs. 4A and B to C, and Fig. 5A to 5B). Thus, apparent limits in bound adjustments in our data under certain conditions (e.g., to trial-by-trial SNR changes shown in Fig. 5C,D) may arise because a wide range of bounds yields near-equivalent reward rates, reducing pressure to adapt. However, changing the incentive structure to sharpen these differences could, in principle, elicit different dynamics. Predictability likely also plays a role: the condition that showed the least adjustment (trial-by-trial SNR changes) was also the least predictable.

These findings also have important implications for the underlying neural mechanisms. Neural correlates of decision bounds have been identified using RT tasks in which a decision variable based on accumulated sensory evidence informs a behavioral response, often a saccadic eye movement to a particular visual target. These decisions are typically assumed to use a fixed bound on the accumulated evidence as the decision rule (Gold and Shadlen, 2007), represented in the brain as a convergence of neural activity to a fixed level at the end of the decision process and triggering the associated action (e.g., by activating subcortical circuits that control saccades (Schall and Thompson, 1999). This kind of “bound-crossing”-like activity has been observed in the FEF of the dorsolateral prefrontal cortex (dlPFC), the lateral intraparietal (LIP) area, premotor cortex, and the superior colliculus in monkeys (Hanes and Schall, 1996 p.199; Roitman and Shadlen, 2002; Ratcliff et al., 2007; Churchland et al., 2008; Ding and Gold, 2012; Thura and Cisek, 2016; Stine et al., 2023). Our results imply that these bound crossings are flexible and depend on both the goals of the decision-maker and the predictability of the factors that are relevant to forming and evaluating the outcome of the decision. More work is needed to better understand how this flexibility is implemented in the brain to allow people (and animals) to find the right balance of speed and accuracy when making deliberative, temporally extended decisions.

In summary, our results show that decision bounds are not set as fixed values applied uniformly across contexts but instead are flexible control variables that depend critically on the temporal structure and predictability of evidence. Using a task that allowed us to infer decision bounds directly on individual trials, we found that people adjust their bounds in qualitatively different, and sometimes opposite, ways depending on whether changes in evidence quality are expected across blocks, across trials, or within a trial. These adjustments are consistent with near-optimal, satisficing strategies shaped by reward-rate structure rather than strict maximization. More broadly, our findings highlight that understanding the flexibility with which we make decisions requires characterizing not only how evidence is accumulated, but also how rules to end the accumulation process and commit to a choice adapt to uncertainty and change.

## Methods

### Task

The pigeon task was built in PsychoPy, converted to Java using built-in PsychoPy features (Peirce et al., 2019), hosted on Pavlovia (pavlovia.org), and run on Prolific (prolific.com). All code (analysis and task deployment) is publicly available at https://github.com/TheGoldLab/Analysis_Pigeon. We recruited three sets of 60 participants (cohort 1 age range=19–34 years, median age=23 years, 29 men, 30 women, 1 not given; cohort 2 age range=19–51 years, median age=23 years, 30 men, 30 women; cohort 3 age range=20–73 years, median age=32.5 years, 35 men, 24 women, one not given).

At the beginning of each session, the participant provided consent and then completed a tutorial that explained the pigeon’s movements, rewards, penalties, and feedback. The tutorial also included a description of the inherent trade-off between speed and accuracy and provided general instructions to be as fast and accurate as possible. The task was then run in blocks of trials, each of which involved a different set of conditions (Table 1).

Across all conditions, a single trial had the same general structure: the pigeon would take a “biased random walk” that started in the center of the screen and then tended to drift towards one of two seed piles located equal distances to the left and right of the center, and at a self-determined time, the participant would push a key to indicate which pile they thought the pigeon would end up at on that trial. Each step of the random walk was governed by taking an independent, identically distributed sample every 200 ms from a Gaussian distribution. This distribution had a standard deviation that was fixed for all trials within the block and a mean that was chosen at random (50/50) from one of two values (a positive number and its negative). Each block was completed when the pigeon took a total of 600 steps across trials. The step count was incremented in three ways: 1) +1 at the beginning of each trial; 2) +1 for each step taken by the pigeon during a trial, before the participant made their choice; and 3) by an additional number of steps applied as an error penalty, when appropriate. Participants were paid a flat fee for finishing all blocks of trials in a given session ($12/hr) and then a bonus based on performance ($1 for each block that they received at least 80% of the total coins that an ideal observer using the optimal bound would achieve).

The three cohorts differed in whether SNR varied across blocks, across trials, or within trials, along with certain costs and rewards. For cohorts 1 and 2, all task parameters were fixed within a trial, including pigeon SNR, defined as the absolute mean of the step distribution divided by its standard deviation, that was either fixed (in absolute value) within a block (all blocks for cohort 1, 3 blocks for cohort two) or selected randomly on each trial within a block (3 blocks for cohort 2). For cohort 3, the first two blocks each used a fixed (high or low) SNR throughout the block. For the third block, the SNR was selected randomly on each trial from the two values (low or high) but was then held fixed throughout the trial. The final two blocks included a SNR changepoint on each trial, one block from low to high SNR and the other from high to low SNR. In both cases, the changepoint occurred at a fixed, cued time within each trial, corresponding to the median (for 40 participants) or mean (for 20 participants) RT for that participant for the starting SNR (i.e., from the low SNR block for low→high changes and from the high SNR block for high→low changes) and indicated via a visual timer and progress bar that counted down to this time.

### Behavioral analysis

Response time (RT) was measured for each trial as the time, in number of pigeon steps from the beginning of the trial, of the keypress. Non-decision time (NDT) was computed separately for each participant and SNR (both within and across blocks) as the number of pigeon steps (restricted to between 0 and 4 based on plausible sensorimotor delays given the step duration) before the keypress that maximized the mean congruence between the pigeon position and the choice across trials (i.e., the fraction of trials in which both were on the same side of the midline). Decision time (DT) was determined for each trial as the RT from that trial minus the NDT from that session. The decision bound was defined for each trial as the distance of the pigeon from the midline to halfway between one step before the DT and at the DT (that is, we assumed that the bound was crossed sometime during this final decision step and took its expected value). Unless otherwise noted, we present the bound as an unsigned quantity representing the absolute distance. This distance has units of normalized horizontal distance from the center of the screen (i.e., the starting pigeon position, where distance=0) relative to the edge of the screen (i.e., distance=1; the seed piles were placed at a distance of 0.8 from the center, which is the maximum point the pigeon could reach).

### Bound corrections

At longer DTs, our method of quantifying bound height (halfway between the final two steps during decision formation) becomes an increasingly unbiased estimator of the true bound (assuming an accumulate-to-bound decision process), because the pigeon step locations become increasingly independent of any chosen bound. However, biases are pronounced at shorter DTs, because the pigeon location is not independent of any chosen bound (e.g., the first step is always distributed across possible bounds according to the pigeon-step generative distribution). Because of this dependence, at short DTs low bounds tend to be overestimated, and high bounds tend to be underestimated (Fig. 7, gray line).

**Figure 7:**
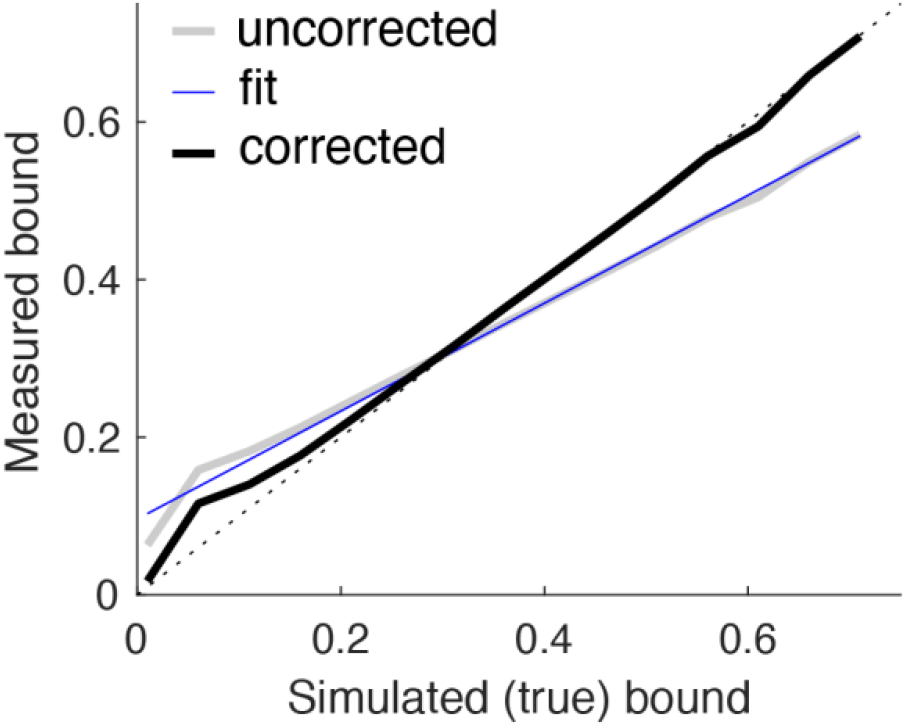
Correcting biases in bound estimates. Bound estimates from pigeon positions were biased in simulations, such that at relatively short DTs low bounds tended to be overestimated, and high bounds tended to be underestimated. We accounted for these biases using linear regressions from simulated data to scale bound estimates. The figure shows data from simulations using a range of bounds but all resulting in choices (bound crossings) at DT=2. The linear fit (blue line) to the uncorrected data (gray line) was used to adjust the corrected data (black line).

To account for these biases, we simulated decisions for each SNR and each of 16 fixed bound values (from 0.01 to 0.76 in steps of 0.05). For these simulations (1,000,000 simulated trials per SNR and bound), we assumed that the NDT on each trial was selected randomly from between 0 and 2 steps (restricting it to just 1–2 did not substantially affect these simulations). Each simulation produced a DT on each trial for the given bound and SNR. For each DT, we combined data across all simulated bounds and compared the “true” (simulated) bound with the measured bound, analyzed as described above. We corrected for biases by fitting linear regressions to these relationships, then used the fits to adjust our measured values such that, as applied to the simulated data, the corrected, measured values equaled the true, simulated values (Fig. 7, black line).

### Reward-rate functions

Reward rate functions of bound height were determined via simulations. Each set of simulations for a given set of task conditions was repeated 500 times for each of 16 bound values (from 0.01 to 0.76 in steps of 0.05). The simulations assumed that on each trial, the bound was either: 1) fixed throughout the trial (i.e., Figs. 4 and 5); or 2) underwent a single changepoint at the time of the SNR changepoint (i.e., Fig. 6).

## Supported by

CRCNS NSF_22-07727 (Josić, Kilpatrick, Ding, and Gold) and R01-EY-034640 (Ding and Gold).

